# The enigmatic fungal genus *Ceraceosorus* provides a theoretical framework for studying intragenomic variation in ribosomal DNA sequences

**DOI:** 10.1101/2024.04.10.588980

**Authors:** Teeratas Kijpornyongpan, Mary Claire Noble, Marcin Piątek, Matthias Lutz, M. Catherine Aime

## Abstract

Multicopy nuclear ribosomal (rDNA) genes have been used as markers for fungal identification for three decades. The rDNA sequences in a genome are thought to be homogeneous due to concerted evolution. However, intragenomic variation of rDNA sequences has recently been observed in many fungi, which cause problems in fungal identification and species abundance estimation. Various sequence-based methods have been used to demonstrate rDNA sequence heterogeneity, but there is no technical assessment of the comparability of results from these methods. In this article, we sampled smut fungi representing all major lineages of subphylum Ustilaginomycotina as a system to examine sequence heterogeneity in the rDNA repeats. Three methods were used: PCR-cloning-Sanger sequencing, targeted amplicon high-throughput sequencing, and WGS high-throughput sequencing. Based on our analyses, *Ceraceosorus* is the only sampled fungal genus in Ustilaginomycotina showing intragenomic variation, with up to 27 nucleotide variant sites in the ITS1–5.8S–ITS2 region and 2.6% divergence among analyzed ITS haplotypes. We found many conflicting patterns across the three detection methods, with up to 28 conflicting variant sites in one sample. Surprisingly, at least 40% of these conflicts are due to PCR-cloning-sequencing errors, as the corresponding variant sites were not observed in the other methods. Based on our data and the literature, we evaluated the characteristics and advantages/disadvantages of each detection method. A model for how intragenomic variation may arise in the rDNA region is presented. Finally, we describe the fourth known species of *Ceraceosorus*, *C. americanus*, isolated from an asymptomatic rosemary leaf collected in Louisiana, USA. We anticipate that our study will provide a framework for future research in rDNA regions as well as other similar multicopy genes.

**Author Summary:** Ribosomal DNA (rDNA) genes are one of the most ancient multicopy genes in cellular organisms. They function as a part of the protein synthesis machinery in a cell. The rDNA sequences have also been used in species identification and microbial community profiling. Despite these utilities, little is known how the rDNA genes have evolved. Biologists initially thought the sequences among rDNA copies are homogeneous, but many recent cases illustrated rDNA sequence heterogeneity. In this article, we utilized the fungal genus *Ceraceosorus* together with allied smut fungi as a system to study sequence heterogeneity in the rDNA genes using various detection methods. Our system found rDNA sequence homogeneity as a common form, while sequence heterogeneity is taxon-specific. Based on our data and literature review, we explained possible sources for sequence heterogeneity in the rDNA genes. Our study also noticed result discrepancies across variant detection methods. These include artefactual variants from the PCR-cloning-sequencing method, inconsistent detected variants from the independent runs of high-throughput sequencing, and technical errors in bioinformatic analyses. We therefore emphasize the importance of methodological choices which have different pros and cons for studying intragenomic variation of rDNA genes, as well as other multicopy gene families.

## Introduction

Nuclear ribosomal DNA (rDNA) genes are among the most conserved and redundant genes in living organisms. They encode ribosomal RNA, which plays a critical role in ribosome assembly and protein synthesis (Bruns et al. 1991; Hillis and Dixon 1991). In typical eukaryotic genomes, there are four types of encoded rRNAs: 18S, which is part of the small subunit of the ribosome, and 5S, 5.8S, and 28S, which together form the large subunit of the ribosome. While the 5S rDNA shows more variability in its presence/absence and its location in the genomes (Drouin and Moniz de Sá 1995), the rDNA sequences encoding 5.8S, 18S, and 28S rRNA are generally clustered as a single operon unit that is tandemly repeated in the genomes. The copy number of rDNA sequences varies among different animal, plant, and fungal species, ranging from a dozen to tens of thousands (Prokopowich et al. 2003; Lofgren et al. 2019). Although there is a positive correlation between genome size and rDNA copy number in animals and plants (Prokopowich et al. 2003), this trend is not observed in fungi (Lofgren et al. 2019).

In contrast to the mutation-selection mechanisms that generally occur within a genome, most rDNA copies in a genome appear to be homogeneous (Hillis and Dixon 1991; Liao 1999; Nei and Rooney 2005; Eickbush and Eickbush 2007). Due to their varying rates of evolution, abundance of copies, sequence homogeneity, and ease of amplification by PCR, the rDNA sequences have served as one of the most powerful markers for taxonomy, systematics, and metagenomics for the past 30 years (Bruns et al. 1991; Kranz et al. 1995; Halanych 2004; Lindahl et al. 2013; Paloi et al. 2022). Sequences from two rDNA regions are currently used as universal DNA barcodes for fungal species identification: the D1/D2 region of 28S rDNA as a DNA barcode for yeasts and the ITS1–5.8S–ITS2 region as a DNA barcode for most fungi (Fell et al. 2000; Schoch et al. 2012).

A phenomenon called “concerted evolution” has been used to explain how most rDNA copies in a genome maintain sequence homogeneity (Elder and Turner 1995; Liao 1999; Eickbush and Eickbush 2007). DNA recombination and gene conversion are two main forces for gene homogenization to transform a variant copy into an original copy, to remove a variant copy from a genome, or to transform an original copy into a variant copy. This mechanism is clearly reviewed and discussed elsewhere (Liao 1999; Eickbush and Eickbush 2007; Paloi et al. 2022). Recombination-mediated gene homogenization has recently been demonstrated in fungal rDNA sequences (Liu et al. 2015; Dakal et al. 2016). However, many recent studies reveal that intragenomic variation of the rDNA sequences is more common throughout the fungal kingdom than previously thought. Hybridization and inefficient concerted evolution are thought to be two major forces for intragenomic rDNA variation in fungi (Paloi et al. 2022). Although intragenomic rDNA variation has been detected by various sequence-based methods (Paloi et al. 2022), a comprehensive technical evaluation of these detection methods has never been performed. In addition, there is no standard convention for the appropriate reporting of intragenomic variation data.

*Ceraceosorus* is an enigmatic fungal genus in Ceraceosorales, Ustilaginomycotina. It was first described as a non-teliosporic plant pathogen, typified by *Ceraceosorus bombacis* from the leaves of *Bombax ceiba* in India (Bakshi et al. 1972; Cunningham et al. 1976), now placed in the Exobasidiomycetes (Ustilaginomycotina) as a sister to species of the Entylomatales (Begerow et al. 2006; Kijpornyongpan et al. 2018). The Entylomatales contains mainly teliosporic smuts that infect dicotyledonous hosts and show a high degree of host specificity (Vánky 2012; Kruse et al. 2018; Piątek et al. 2024). The genus *Ceraceosorus* currently consists of three described species: *C. africanus, C. bombacis*, and *C. guamensis* (Cunningham et al. 1976; Kijpornyongpan and Aime 2016; Piątek et al. 2016). A yeast-like asexual stage has been described only in *C. guamensis*, while the other two species are known only from the sexual stage as phytopathogens on the leaves of *Bombax* spp. Our previous study revealed extensive intragenomic variation in the internal transcribed spacer 1–5.8S–internal transcribed spacer 2 (ITS) region of rDNA repeats in *C. bombacis* and *C. guamensis* (Kijpornyongpan and Aime 2016). In the present study, we describe a fourth species, *C. americanus*, which is the first report of the genus in the Americas. We then investigate intragenomic rDNA variation in all four species of *Ceraceosorus* and from representatives of additional lineages in Ustilaginomycotina. The study includes the use of three sequence-based methods including PCR-cloning-Sanger sequencing, targeted amplicon high-throughput sequencing, and whole genome shotgun (WGS) high-throughput sequencing. We then compare detected variants across the three methods to determine consensus variant sites, to cross-check for data consistency, and to determine technical challenges for each detection method. Finally, a model for the origin of rDNA sequence heterogeneity is proposed and discussed.

## Materials and Methods

### Sample used for this study

The fungal strain SA252w was isolated from an abaxial leaf surface of *Rosmarinus officinalis* (Lamiaceae) collected on the campus of Louisiana State University, Baton Rouge, Louisiana, USA. Isolation followed the spore drop method described in Albu et al. (2015). The culture was stored on potato dextrose agar (PDA) slants at 4 °C and in 15% glycerol at -80 °C. A dried culture plate was vouchered in the Kriebel Herbarium, Purdue University (PUL) under the voucher number F22392. An ex-type culture has been deposited in the Agricultural Research Service (USDA-ARS) Culture Collection and the Westerdijk Fungal Biodiversity Institute Culture Collection (CBS) under accession nos. NRRL 66778 and CBS 144261, respectively.

Other *Ceraceosorus* strains/specimens included in this study were as follows: the culture collection nos. ATCC 22867 and MCA4658 (≡ CBS 139631 ≡ NRRL 66309) for the ex-type strains of *C. bombacis* (from the leaves of *Bombax ceiba* in India) and *C. guamensis* (from the surface of a healthy dicot leaf in Guam), respectively, and KRAM F-57385, KRAM F-57386, and KRAM F-57387 for the vouchered specimens of *C. africanus* (from the leaves of *Bombax costatum* in Benin and Ghana), deposited at the fungal collection of the W. Szafer Institute of Botany, Polish Academy of Sciences, Kraków.

We also included several other species representing different lineages in Ustilaginomycotina: *Testicularia cyperi* MCA3645 (≡ ATCC MYA-4640) and *Mycosarcoma maydis* (syn. *Ustilago maydis*) TKC58 (≡ NRRL Y-64004) for Ustilaginales, *Meira miltonrushii* MCA3882 (≡ CBS 12591) and *Laurobasidium hachijoense* (syn. *Acaromyces ingoldii*) MCA4198 (≡ CBS 140884) for Exobasidiales, *Tilletiopsis washingtonensis* MCA4186 (≡ NRRL Y-63783) for Entylomatales, *Pseudomicrostroma glucosiphilum* MCA4718 (≡ CBS 14053) and *Jaminaea rosea* MCA5214 (≡ CBS 14051) for Microstromatales, *Violaceomyces palustris* SA807 (≡ CBS 139708) for Violaceomycetales, and *Tilletiaria anomala* CBS 436.72 for Georgefischeriales.

### Culture description

To observe colony morphology, the strain SA252w was cultured on four types of media: PDA, corn meal agar (CMA), yeast malt agar (YMA), and yeast peptone glucose agar (YPGA). The cultures were incubated at room temperature (25 °C) for 14 days. Colony color was compared to color codes from The Online Auction Color Chart^TM^ (Online Auction Color Chart 2004). For cell morphology, the strain was inoculated in yeast malt broth (YMB) and incubated for 7 days at room temperature and 200 round-per-minute shaking condition. A drop of the culture suspension was then placed on a microscope slide, followed by a coverslip. We observed the cell morphologies using the Olympus BH2 microscopy in differential interference contrast mode. Photomicrographs were captured by QImaging with QI Imaging software. Cell sizes were measured for a minimum of 30 cells. The physiological profiles of SA252w were examined using a protocol described in a previous study (Kijpornyongpan and Aime 2016).

### DNA extraction, rDNA amplification and sequencing

DNA was extracted from fresh cultures growing on PDA at room temperature. A piece of colony was cut for extraction using the Omega fungal DNA HP kit (Norcross, GA, USA) following the manufacturer’s protocol. For the vouchered specimens of *C. africanus,* DNA was extracted as described in Piątek et al. (2016).

To amplify sequences in rDNA repeats, a list of primers was used to amplify different rDNA regions: LR0R/LR6 primers (Vilgalys and Hester 1990; Vilgalys et al. 1994) for partial 28S large subunit rDNA, PNS1/NS6 primers (White et al. 1990; Hibbett 1996) for partial 18S small subunit rDNA. ITS1-5.8S-ITS2 region, we used ITS1Cb (5’- CTT GCT GGC CCG GAG GAA GTA A-3’, designed in this study) and ITS1F (Gardes and Bruns 1993) as forward primers for *Ceraceosorus* and other fungal genera, respectively, and ITS4 (White et al. 1990) as a reverse primer for all studied species. Amplification conditions were as described in previous studies (Kijpornyongpan and Aime 2016; Kijpornyongpan and Aime 2017). PCR products were run on 1% agarose gel electrophoresis to determine size and quality.

The rDNA amplicons were sequenced using three strategies. The total PCR products were sent to GENEWIZ (Plainfield, NJ, USA) for direct Sanger sequencing. The ITS region of *C. africanus* and *Ceraceosorus* sp. SA252w could not be directly sequenced due to intragenomic variation among the separated copies. We therefore performed cloning-Sanger sequencing as described in the previous study (Kijpornyongpan and Aime 2016). At least 10 clones were sequenced for each strain/specimen. For targeted amplicon sequencing, which allows for sequence heterogeneity and provides greater coverage of sequenced regions, PCR products from all rDNA regions for each species were pooled into a single sample and sent to the Purdue Genomics Core Facility for Illumina MiSeq sequencing. Library preparation and sequencing were performed using Illumina NexteraXT (Illumina, San Diego, CA, USA) and Illumina MiSeq 500 cycle kits, respectively. About 50,000 – 200,000 Illumina reads were generated for each sample. Finally, adapter sequences and low-quality bases were trimmed using Cutadapt 1.12 (Martin 2011) and Trimmomatics 0.36 (Bolger et al. 2014).

### Phylogenetic analyses

To determine the phylogenetic placement of the strain SA252w, 18S, 28S and ITS sequences from all described *Ceraceosorus* species and representative fungi of Ustilaginomycotina (see above) were retrieved from the literature and otherwise sequenced in this study (Table 1). *Mixia osmundae* was selected as an outgroup based on its placement in Pucciniomycotina, a sister subphylum to Ustilaginomycotina (Prasanna et al. 2019). Sequences of each rDNA region were aligned using the MUSCLE algorithm performed in MEGA-X (Kumar et al. 2018). Flanking alignment regions at 5’ and 3’ were manually trimmed. The trimmed alignments of all three rDNA regions were then concatenated and used for phylogenetic reconstruction, which was performed through RAxML 8.2.9 (Stamatakis 2014) with GTRCAT as a nucleotide substitution model and 1000-replicate bootstrapping.

**Table 1.**
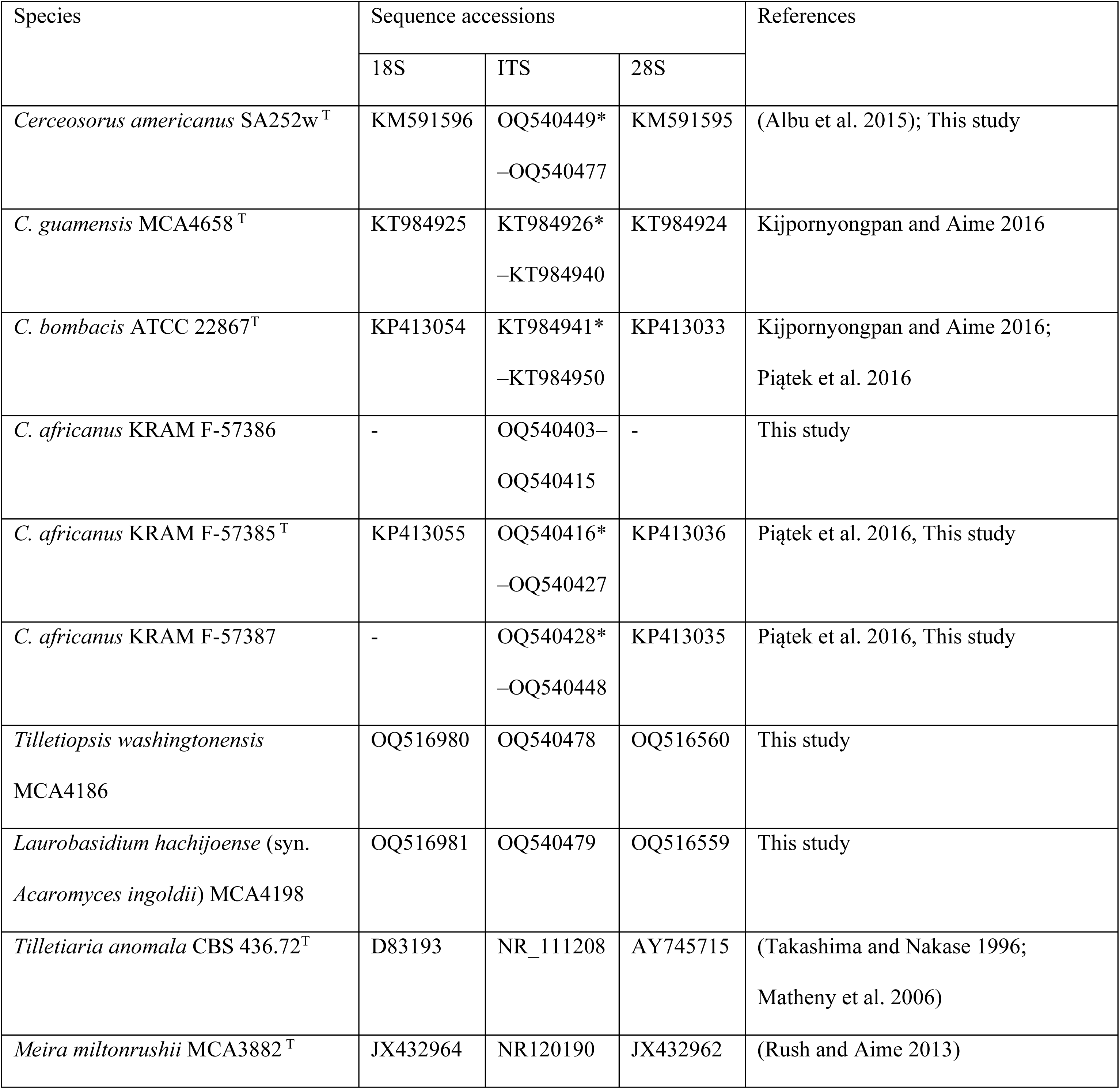
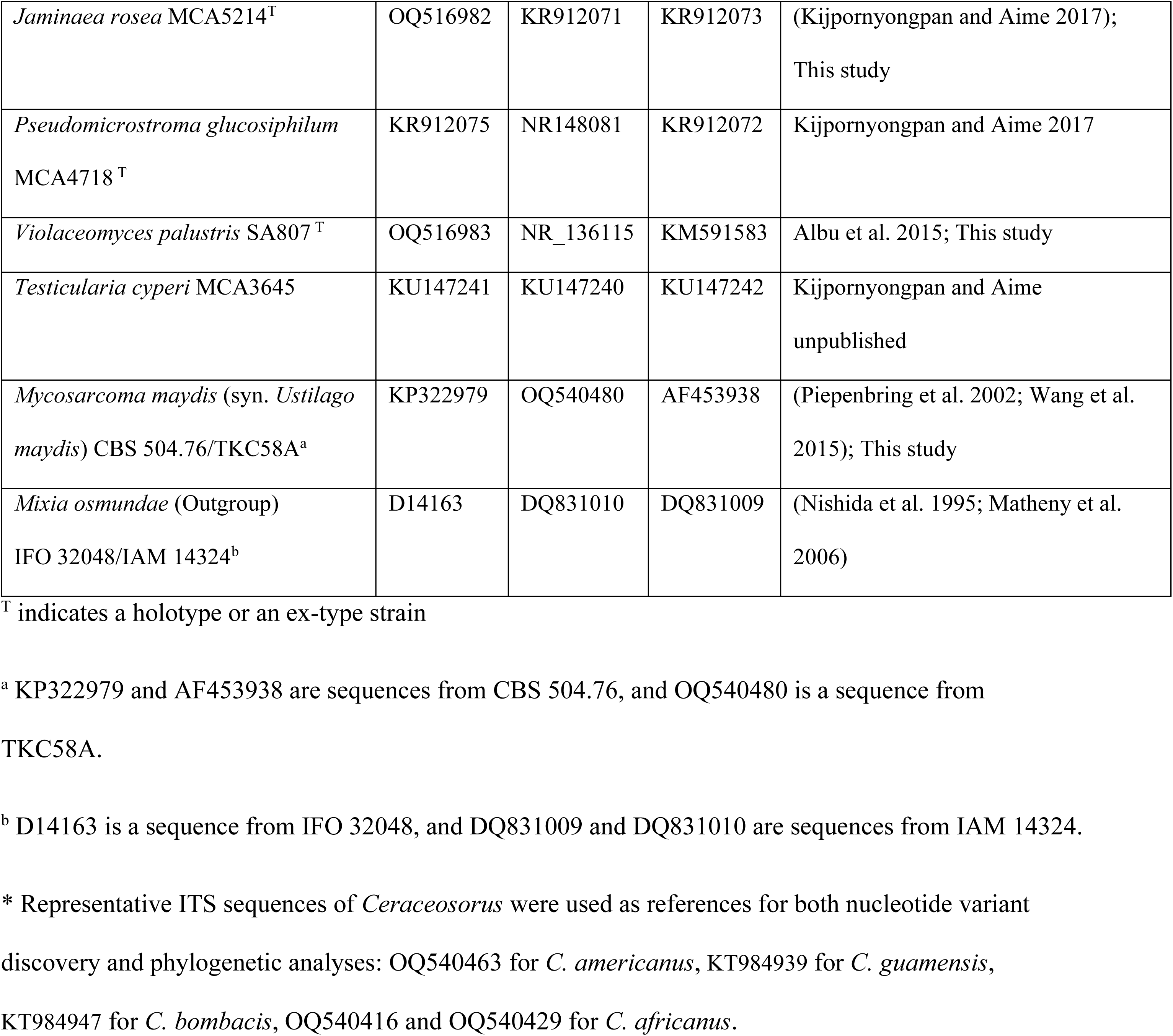
rDNA sequences used in this study.

| Species | Sequence accessions |  |  | References |
| --- | --- | --- | --- | --- |
|  | 18S | ITS | 28S |  |
| *Cerceosorus americanus* SA252w<sup>T</sup> | KM591596 | OQ540449*<br>–OQ540477 | KM591595 | (Albu et al. 2015); This study |
| *C. guamensis* MCA4658<sup>T</sup> | KT984925 | KT984926*<br>–KT984940 | KT984924 | Kijpornyongpan and Aime 2016 |
| *C. bombacis* ATCC 22867<sup>T</sup> | KP413054 | KT984941*<br>–KT984950 | KP413033 | Kijpornyongpan and Aime 2016;<br>Piątek et al. 2016 |
| *C. africanus* KRAM F-57386 | - | OQ540403–<br>OQ540415 | - | This study |
| *C. africanus* KRAM F-57385<sup>T</sup> | KP413055 | OQ540416*<br>–OQ540427 | KP413036 | Piątek et al. 2016, This study |
| *C. africanus* KRAM F-57387 | - | OQ540428*<br>–OQ540448 | KP413035 | Piątek et al. 2016, This study |
| *Tilletiopsis washingtonensis*<br>MCA4186 | OQ516980 | OQ540478 | OQ516560 | This study |
| *Laurobasidium hachijoense* (syn.<br>*Acaromyces ingoldii*) MCA4198 | OQ516981 | OQ540479 | OQ516559 | This study |
| *Tilletiaria anomala* CBS 436.72<sup>T</sup> | D83193 | NR_111208 | AY745715 | (Takashima and Nakase 1996;<br>Matheny et al. 2006) |
| *Meira miltonrushii* MCA3882<sup>T</sup> | JX432964 | NR120190 | JX432962 | (Rush and Aime 2013) |
| *Jaminalia rosea* MCA5214<sup>T</sup> | OQ516982 | KR912071 | KR912073 | (Kijpornyongpan and Aime 2017);<br>This study |
| *Pseudomicrostroma glucosiphilum*<br>MCA4718<sup>T</sup> | KR912075 | NR148081 | KR912072 | Kijpornyongpan and Aime 2017 |
| *Violaceomyces palustris* SA807<sup>T</sup> | OQ516983 | NR_136115 | KM591583 | Albu et al. 2015; This study |
| *Testicularia cyperi* MCA3645 | KU147241 | KU147240 | KU147242 | Kijpornyongpan and Aime<br>unpublished |
| *Mycosarcoma maydis* (syn. *Ustilago*<br>*maydis*) CBS 504.76/TKC58A<sup>a</sup> | KP322979 | OQ540480 | AF453938 | (Piepenbring et al. 2002; Wang et al.<br>2015); This study |
| *Mixia osmundae* (Outgroup)<br>IFO 32048/IAM 14324<sup>b</sup> | D14163 | DQ831010 | DQ831009 | (Nishida et al. 1995; Matheny et al.<br>2006) |

<sup>T</sup> indicates a holotype or an ex-type strain
<sup>a</sup> KP322979 and AF453938 are sequences from CBS 504.76, and OQ540480 is a sequence from TKC58A.
<sup>b</sup> D14163 is a sequence from IFO 32048, and DQ831009 and DQ831010 are sequences from IAM 14324.
\* Representative ITS sequences of *Ceraceosorus* were used as references for both nucleotide variant discovery and phylogenetic analyses: OQ540463 for *C. americanus*, KT984939 for *C. guamensis*, KT984947 for *C. bombacis*, OQ540416 and OQ540429 for *C. africanus*.

To analyze the sequence variation of *Ceraceosorus* ITS regions, all sequenced ITS amplicons from all *Ceraceosorus* species were aligned in MEGA-X using the MUSCLE algorithm. The alignment was used for p-distance calculation using the MEGA-X built-in function, and for phylogenetic reconstruction using the Neighbor-Joining (NJ) method performed in MEGA-X with the Kimura-2 parameter as a nucleotide substitution model and 1000-replicate bootstrapping.

### Intragenomic nucleotide variant detection

We detected intragenomic nucleotide variants within rDNA regions (partial 18S, partial 28S and ITS) from three data types—cloning-Sanger sequencing, targeted amplicon Illumina sequencing, and whole genome shotgun (WGS) Illumina sequencing. For targeted amplicon sequencing, the cleaned Illumina MiSeq reads from *Ceraceosorus* species, *M. maydis* (syn. *U. maydis*), *M. miltonrushii*, *P. glucosiphilum*, *T. washingtonensis*, and *V. palustris* were mapped to reference rDNA sequences (Table 1) using Bowtie 2 2.3.3.1 (Langmead and Salzberg 2012), followed by conversion to BAM alignment files using Samtools 1.6 (Li et al. 2009). Variant detection was then performed with GATK 3.8.0 (McKenna et al. 2010) using the HaploTypeCaller and DeptOfCoverage functions with default parameters.

For WGS Illumina sequences, cleaned reads from the WGS sequencing projects of several Ustilaginomycotina species were used for nucleotide variant detection: *C. guamensis*, *J. rosea*, *L. hachijoense* (syn. *A. ingoldii*), *M. miltonrushii*, *P. glucosiphilum*, *Testicularia cyperi*, *Tilletiaria anomala*, *T. washingtonensis*, and *V. palustris* (Toome et al. 2014; Kijpornyongpan et al. 2018). Reads were retrieved from the sequencing read archive of the JGI MycoCosm genome portal (Grigoriev et al. 2014). For *C. bombacis*, the 300-bp paired-end WGS sequencing reads from a previous study (Sharma et al. 2015), under the accession ERX541976 (the run accession ERR583964), were downloaded from the NCBI Sequence Read Archive database through the sra-toolkit. The variant detection was performed using the procedure described above. We also estimated the rDNA copy number of each genome by a following calculation: a percentage of reads mapped to reference rDNA sequences x genome assembly size (in bp) / length of reference rDNA sequences (in bp) / a percentage of reads mapped to genome assembly. Any call with fewer than 500 mapped reads and a variant frequency of less than 0.10 was discarded from the valid list of nucleotide variants, unless the variants were present in at least two independent runs of Illumina sequencing (including targeted amplicon sequencing and WGS sequencing).

For the ITS sequences of *Ceraceosorus* species obtained from PCR-cloning-Sanger sequencing, we aligned the ITS sequences of each species using the MUSCLE algorithm run in MEGA-X. The variant detection was done by visual inspection of the alignments. Any site having a variant site present in only one sequence for each strain was excluded from the valid variant list and we considered it as a PCR-cloning-sequencing error, unless the variant site was also present in other detection methods. Finally, the valid nucleotide variants were used to correct PCR-cloning-sequencing errors from the *Ceraceosorus* ITS alignment. The corrected ITS alignment was subjected to another p-distance calculation and phylogenetic reconstruction to evaluate the effect of PCR-cloning-sequencing errors. Overview of our analysis pipeline in study can be found in Figure 1.

**Figure 1.**
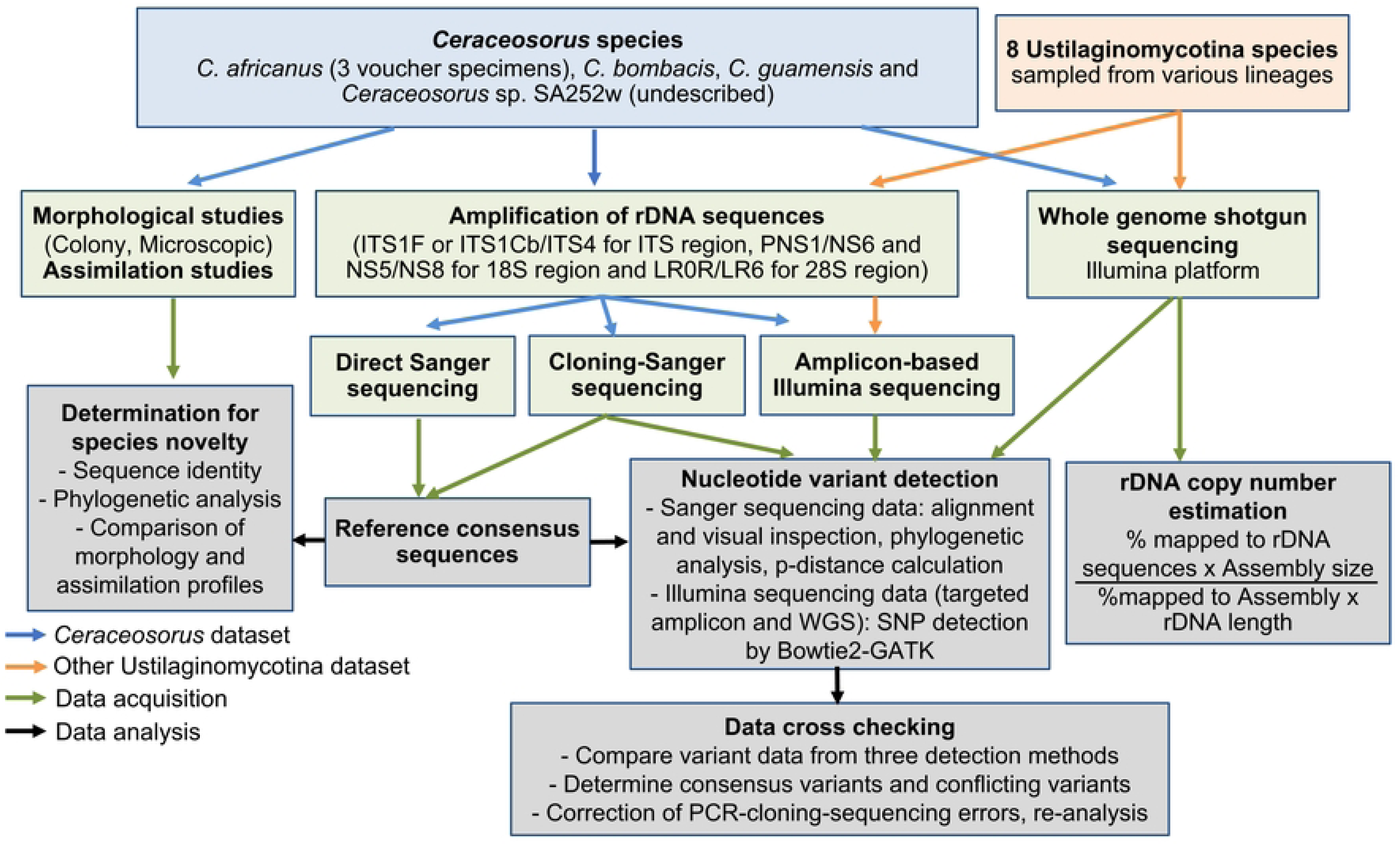
Analysis pipeline for this study.

### Data visualization

All phylogenies were visualized in MEGA-X. We used the ggplot2 package (Wickham 2016), run in R 3.5.2 and RStudio 1.4.1103 to visualize detected nucleotide variants in the *Ceraceosorus* ITS region. Other graphics were created and edited in Inkscape 0.92 (https://www.inkscape.org).

## Results

### Taxonomy

*Ceraceosorus americanus* T. Kij. & Aime sp. nov. (Figure 2, Table 2)

**Figure 2.**
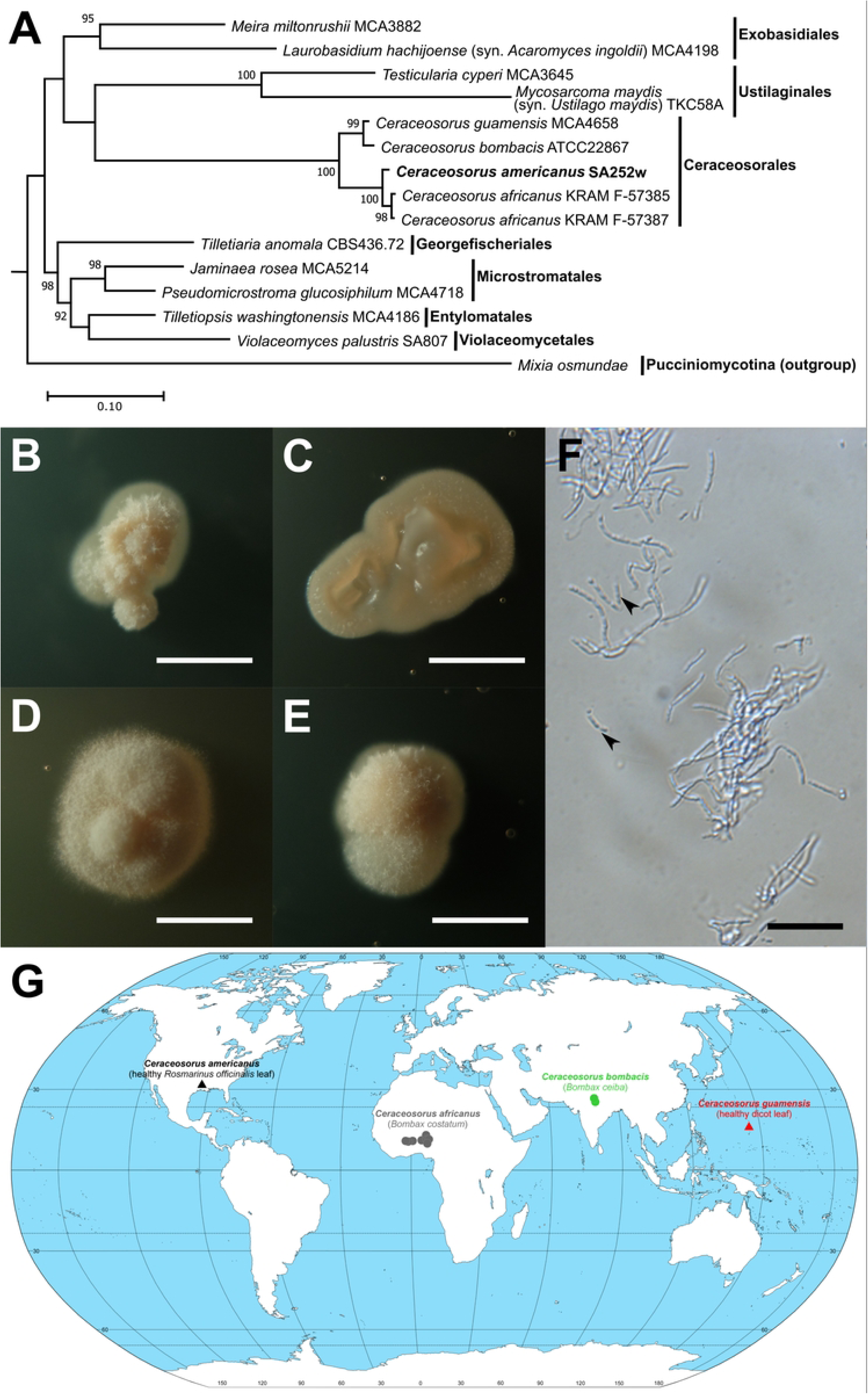
*Ceraceosorus americanus* sp. nov. SA252w **(A**) A phylogenetic tree confirming the placement of the strain SA252w in Ceraceosorales (Ustilaginomycotina). The rDNA sequences (Table 1) of representative Ustilaginomycotina species and all described *Ceraceosorus* species, were aligned and concatenated. The alignment was used for phylogenetic reconstruction using RAxML with GTRCAT as the nucleotide substitution model and 1000-replicate bootstrapping. Bootstrap values are only shown for nodes with strong support (> 70%). *Mixia osmundae* was selected as an outgroup based on its placement in a different subphylum (Pucciniomycotina). **(B) – (E)** Colony morphology of SA252w on potato dextrose agar (B), corn meal agar (C), yeast malt agar (D), and yeast peptone glucose agar (E). Bars: 2.5 mm. **(F)** Cell morphology of SA252w cultured in yeast malt broth (YMB) for 14 days. Cell suspension observed under a DIC microscope. Arrowheads indicate conidia. Bar: 20 μm. **(G)** Worldwide distribution of *Ceraceosorus* species. Sources of isolation are indicated in parentheses. Dots indicate phytopathogenic species with sexual stages. Triangles indicate species known only from asexual stages.

**Table 2.** Compounds that have different assimilation profiles among *Ceraceosorus* ex-type strains: ATCC 22867 for *Ceraceosorus bombacis*, MCA4658 for *Ceraceosorus guamensis*, and SA252w for *Ceraceosorus americanus.* All *Ceraceosorus* strains can assimilate these compounds in common: D-glucose, sucrose, maltose, maltotriose, D-melezitose, D-raffinose, D-stachyose, L-arabinose, trehalose, turanose, D-mannitol, D-arabitol, *i*-erythritol, galactitol, succinic acid and soluble starch for carbon assimilation, and NaNO_2_ and KNO_3_ for nitrogen assimilation. They can also grow in vitamin-free medium, as well as in two osmotic media: 50% glucose YE agar and 10& NaCl YE agar. The maximum temperature all strains can grow in liquid culture is 30 °C. None of these strains assimilates these compounds: D-psicose, L-rhamnose, L-sorbose, α-methyl D-glucoside, D-galacturonate, D-glucuronate, methyl succinate, citrate, inositol, arbutin and salicin for carbon assimilation and L-lysine, cadaverine, creatine and creatinine for nitrogen assimilation.

| Assimilation | ATCC 22867 | MCA4658 | SA252w |
| --- | --- | --- | --- |
| **Carbon compound** |  |  |  |
| Cellobiose | - | - | + |
| Lactose | - | - | + |
| D-Galactose | - | - | + |
| Palatinose | + | - | + |
| D-Melibiose | + | w | + |
| D-Arabinose | - | + | + |
| D-Ribose | w | + | + |
| Glycerol | - | + | + |
| D-Xylose | - | + | w |
| Gentiobiose | - | + | + |
| Ethanol | - | - | + |
| Maltitol | - | w | s |
| Fumaric acid | w | + | - |
| Xylitol | s | + | + |
| 1,2-propanediol | - | - | + |
| Lactate | - | + | - |
| Methanol | w | - | + |
| **Nitrogen compound** |  |  |  |
| Ethylamine-HCl | w | + | - |

Sign: +, growth; -, no growth; w, weak growth; s, slow growth

MycoBank accession number: MBXXXXX (TO BE ADDED UPON THE ACCEPTANCE FOR PUBLICATION)

Holotype: USA. Louisiana, East Baton Rouge Parish. Louisiana State University Baton Rouge Campus. Isolated from the leaf surface of *Rosmarinus officinalis* (Lamiaceae) using the spore drop method. 17 February 2011, leg. S. Albu; PUL F22392, holotype as a dried culture; SA252w ≡ NRRL 66778 ≡ CBS 144261, ex-type cultures; GenBank accessions: KM591596, 18S ribosomal DNA sequence; KM591595, 28S ribosomal DNA sequence; OQ540449–OQ540477, ITS1-5.8S-ITS2 ribosomal DNA sequence.

Etymology: *americanus*, referring to the continent where the fungus was isolated, as this is the first species of *Ceraceosorus* found and collected in the Americas.

Description: Colony morphology. After 14 days of incubation at room temperature (25 °C), the colony morphologies of *C. americanus* are as follows (Figure 2B – E). Colonies on PDA are 2.5 mm in diameter, light cream (oac815), dull, velvety with filiform margins and elevated folds. Colonies on CMA are 3.5 mm in diameter, light cream (oac857), glabrous, butyrous with entire margins and flat elevations. Colonies on YMA are 4 mm in diameter, white cream (oac795), dull, velvety with filiform margins and elevated folds. Colonies on YPGA are 3 mm in diameter, light cream (oac814), dull, velvety with filiform margins and convex (dome) elevation. The average colony growth rate is 3.5 mm per 14 days, with the highest growth rate when cultured on YMA. No sexual reproduction was observed.

Micromorphology. After incubation in YMB with 200 round-per-minute shaking condition at room temperature (25 °C) for 7 days, the cell colony morphologies of *C. americanus* are as follows (Figure 2F). Filamentous growth appears as true hyphae with diameters of 1.1 – 1.7 mm. Cell lengths in the hyphae vary from 1.8 to 6.8 mm. Hyphae are branched and septate. No clamp connection is formed. Conidia appear as one or a few cells detached from the tip of the hyphae.

Physiological properties. *Ceraceosorus americanus* can assimilate the following compounds as a carbon source: D-glucose, D-galactose, sucrose, cellobiose, lactose, maltose, maltotriose, D-melezitose, D-melibiose, D-raffinose, D-stachyose, D-ribose, D-arabinose, L-arabinose, D-xylose (weak), gentiobiose, palatinose, trehalose, turanose, D-mannitol, D-arabitol, *i-*erythritol, galactitol, maltitol (slow), xylitol, methanol, ethanol, 1,2-propanediol, succinic acid, and soluble starch. The fungus can assimilate potassium nitrate (KNO_3_) and sodium nitrite (NaNO_2_) as nitrogen sources. The following compounds are not assimilated by *C. americanus*: D-psicose, L-rhamnose, L-sorbose, α-methyl D-glucoside, D-galacturonate, D-glucuronate, methyl succinate, citrate, fumarate, lactate, inositol, arbutin, and salicin for carbon assimilation and ethylamine, L-lysine, cadaverine, creatine and creatinine for nitrogen assimilation. The fungus can grow in vitamin-free medium, as well as in two hyperosmotic media: 50% glucose yeast extract agar and 10% sodium chloride yeast extract agar. Fermentation is absent. The maximum tested temperature at which the fungus can grow is 30 °C.

Notes: *Ceraceosorus americanus* is the first *Ceraceosorus* species described from the Americas, while the other species are known from the Old World tropics: *C. africanus* from West Africa, *C. bombacis* from India, and *C. guamensis* from Guam (Figure 2G). *Ceraceosorus africanus* and *C. bombacis* are known only from the sexual morph as plant pathogens on the leaves of *Bombax costatum* and *B. ceiba*, respectively. However, *C. americanus* and *C. guamensis* were both isolated from asymptomatic leaf phylloplanes and are only known from the asexual morph; it is unknown whether they also have a phytopathogenic strategy and sexual stages in their life cycle. The two phytopathogenic species live on *Bombax* hosts, which may indicate host specialization of the genus *Ceraceosorus* on this plant genus. In this case *C. americanus* and *C. guamensis* may have lost the phytopathogenic sexual stage because the potential hosts of the genus *Bombax* are absent in the Americas and Guam.

*Ceraceosorus americanus* differs from the other described *Ceraceosorus* species in a few ways. The physiological profiles show several compounds that are assimilated differently among *C. americanus*, *C. bombacis*, and *C. guamensis* (Table 2). Phylogenetic reconstruction of rDNA sequences indicates a distinct placement of *C. americanus* from the other species (Figure 2A). Distinct clustering of ITS clone sequences also confirms the species identity of *C. americanus* (Figure S1).

### Analyses of intragenomic variation in the ITS regions of Ceraceosorus

First, we calculated a p-distance matrix to determine the degree of sequence heterogeneity in the *Ceraceosorus* ITS clone samples (Table S1). Maximum pairwise p-distances of ITS clones observed in each strain/specimen are as follows: 0.031 for *C. bombacis* ATCC 22867, 0.029 for *C. guamensis* MCA4658, 0.019 for *C. americanus* SA252w, 0.025 for *C. africanus* KRAM F-57386, and 0.029 for *C. africanus* KRAM F-57385 and KRAM F-57387. When all three specimens are considered, the maximum pairwise p-distance found in *C. africanus* is increased to 0.035, which exceeds the conventional 97% ITS identity cutoff used to separate OTUs in various studies (Tedersoo et al. 2015).

Phylogenetic reconstruction of *Ceraceosorus* ITS clones reveals that the intragenomic variation is intraspecific—no disparate ITS clone that overlaps across species is found (Figures 3A, S1). We found variability among the three isolates of *C. africanus,* as the ITS clone clustering pattern is not random. The ITS clones of *C. africanus* KRAM F-57386 have less intragenomic variation (average p-distance 0.012) than KRAM F-57385 (average p-distance 0.015) and KRAM F-57387 (average p-distance 0.014). The neighbor-joining tree also indicates isolated placements of the KRAM F-57386 ITS clones compared to the ITS clones from the other two specimens. This correlates to the distant geographic location of KRAM F-57386 (collected from Benin) compared to KRAM F-57385 and KRAM F-57387 (collected from Ghana).

**Figure 3.**
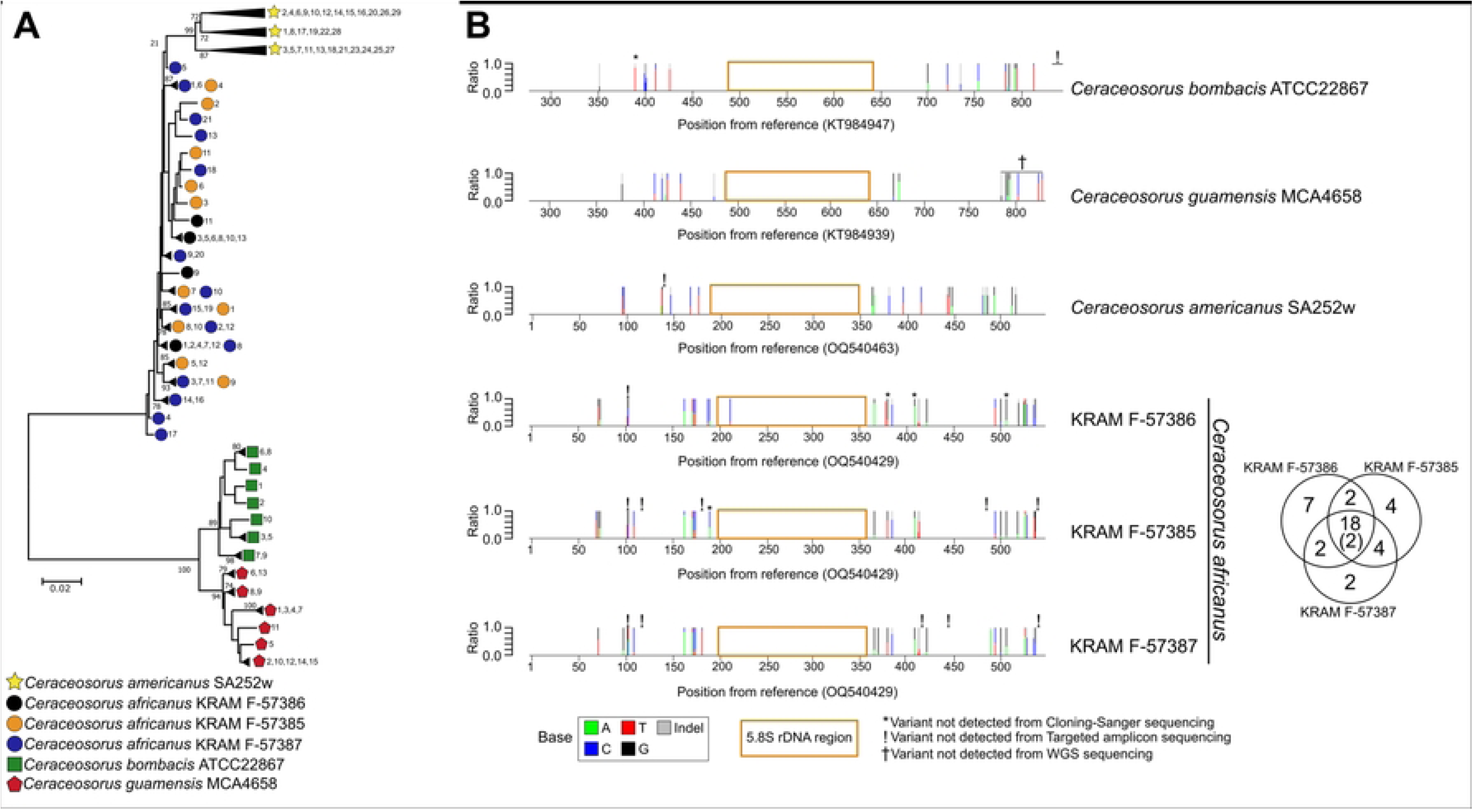
Intragenomic nucleotide variant detection of the internal transcribed spacer (ITS) regions among *Ceraceosorus* species. **(A)** The neighbor-joining tree of the ITS clones of all *Ceraceosorus* samples. The ITS amplicons from the PCR were cloned using T-vector and sequenced using the Sanger platform. The sequenced ITS regions were then aligned and used for neighbor-joining tree reconstruction. For better visualization, all sequences with branch lengths less than 0.01 substitutions/site are collapsed as a subtree. Only bootstrap values greater than 50 are shown on nodes. A symbol in each tip indicates a strain/specimen followed by clone number(s), which is in numerical order from a list of ITS clones (Table 1). The detailed phylogenetic tree can be found in Figure S1. **(B)** Mapping of nucleotide variants in the ITS regions of all *Ceraceosorus* samples. The ITS amplicons from the PCR were used for targeted amplicon Illumina sequencing. Illumina raw reads were then mapped to the reference ITS sequences for nucleotide variant discovery. For each stacked barplot, the X-axis is a position (in base pair) of the reference ITS sequence. The Y-axis is the ratio of nucleotide variants (A, T, C, G or indels) for each position. A position without a stacked line implies no heterogeneity (i.e., no intragenomic variation). Symbols over stacked lines indicate variants that are not detected in different discovery methods: cloning-Sanger sequencing, targeted amplicon sequencing, and WGS sequencing. A gold-framed rectangle on each plot indicates the 5.8S rDNA region of the reference sequence. A Venn diagram at the right side of *C. africanus* tabs indicates numbers of shared variant sites across three samples. Parentheses indicate numbers of shared variant sites in three samples but have conflicts in reference/variant nucleotides. Details of nucleotide variants can be found in Supplementary Table S2.

Our nucleotide variant detection showed that there are at least 15 variant sites in the ITS regions for each *Ceraceosorus* species—no variant site overlaps across different species (Figure 3B, Tables 3, S2). For *C. africanus*, we see common variant sites in three specimens, and most common sites have consistent variant patterns (reference/variant base). However, KRAM F-57386 has more unique variant sites compared to the other two samples, 7 sites compared to 4 and 2 sites, respectively (Figure 3B). One of the unique variant sites is in the 5.8S rDNA region. The reference ITS positions 500 and 537 are two common variant sites that have conflicting patterns—KRAM F-57385 shows different variant bases compared to the other two samples.

**Table 3.** Summary of intragenomic nucleotide variants of *Ceraceosorus* species and other Ustilaginomycotina species detected by targeted amplicon while genome shotgun (WGS) Illumina sequencing.

| Species<br>Length of reference rDNA sequences<br>(partial 18S, ITS, partial 28S) | Number of discovered variant sites in each rDNA<br>region (substitutions/indels) |  |  |  |  |  | Estimated rDNA<br>copy number from<br>WGS data |
| --- | --- | --- | --- | --- | --- | --- | --- |
|  | Targeted amplicon-Illumina sequencing |  |  | WGS-Illumina<br>sequencing |  |  |  |
|  | 18S | ITS* | 28S | 18S | ITS | 28S |  |
| *Ceraceosorus bombacis* ATCC 22867<br>(1683, 846, 2165) | 5/0 | 9/8<br>(12/9) | 5/1 | 5/0 | 12/10 | 8/1 | 8 |
| *Ceraceosorus guamensis* MCA4658<br>(1701, 849, 919) | 3/1 | 11/5<br>(12/5) | 9/0 | 3/1 | 6/4 | 9/0 | 7 |
| *Ceraceosorus africanus* KRAM F-57386<br>(715, 592, 1054) | 9/1 | 18/9<br>(15/8) | 4/0 | NA | NA | NA | NA |
| *Ceraceosorus africanus* KRAM F-57385<br>(715, 592, 1054) | 6/1 | 21/5<br>(20/6) | 0/0 | NA | NA | NA | NA |
| *Ceraceosorus africanus* KRAM F-57387<br>(715, 592, 1054) | 5/1 | 18/7<br>(21/7) | 2/0 | NA | NA | NA | NA |
| *Ceraceosorus americanus* SA252w<br>(1052, 592, 1347) | 2/0 | 13/8<br>(14/8) | 3/2 | NA | NA | NA | NA |
| *Meira miltonrushii* MCA3882<br>(1356, 614, 1076) | 0/1 | 0/0 | 0/0 | 0/0 | 0/0 | 0/0 | 14 |
| *Laurobasidium hachijoense* (syn. *Acaromyces ingoldii*) MCA4198<br>(1713, 573, 1025) | NA | NA | NA | 0/0 | 0/0 | 0/0 | 16 |
| *Tilletiaria anomala* UBC591<br>(1749, 625, 1385) | NA | NA | NA | 0/0 | 0/0 | 0/0 | 233 |
| *Tilletiopsis washingtonensis* MCA4186<br>(1720, 642, 1025) | 1/0 | 0/0 | 0/0 | 0/0 | 0/0 | 0/0 | 30 |
| *Pseudomicrostroma glucosiphilum*<br>MCA4718 (1644, 688, 983) | 0/0 | 1/0 | 0/0 | 0/0 | 1/0 | 0/0 | 17 |
| *Jaminalia rosea* MCA5214<br>(1618, 620, 1019) | NA | NA | NA | 0/0 | 0/0 | 0/0 | 25 |
| *Violaceomyces palustris* SA807<br>(1306, 676, 1037) | 0/0 | 0/0 | 0/0 | 0/0 | 0/0 | 0/0 | 114 |
| *Mycosarcoma maydis* (syn. *Ustilago maydis*) TKC58A<br>(1954, 753, 1115) | 0/0 | 0/0 | 0/0 | NA | NA | NA | NA |

\*Parentheses indicate numbers of variant sites for substitutions/indels detected by cloning-Sanger
sequencing method.
NA: not available, no data, analysis not conducted

We noticed conflicts of observed variant sites in three detection methods (Figure 3B, Tables 4, S2). There are several variant sites that are detected by targeted amplicon Illumina sequencing but not by the PCR-cloning-Sanger sequencing—this is likely due to inadequate sampling efforts. Many observed variant sites from PCR-cloning-Sanger sequencing, with only one observed clone, are likely due to PCR-cloning-sequencing errors, as they are not detected by targeted amplicon Illumina sequencing and WGS sequencing (Tables 4, S2). This is the most common source of conflict in variant calling. We also detected variant sites uniquely found in PCR-cloning-Sanger sequencing with two or more observed clones. Some of these sites are located in the 3’ end of ITS amplicons, making the nucleotide variants barely detectable in the targeted amplicon Illumina sequencing (Figure 3B, Tables 4, S2). Some other sites are undetected, likely due to errors in the high-throughput sequencing, read mapping, or bioinformatic pipeline. On the other hand, these errors can lead to false variant calls in the targeted amplicon Illumina sequencing (Tables S2), which are determined by low coverage (less than 500 read depth), low variant frequencies (less than 0.10), and inconsistency of variant detection between independent runs (approximately 6 – 20% of observed variant sites in each sample have inconsistent detection). Finally, incorrect variant calls can result from inaccurate reference sequences. Thus, an observed nucleotide variant site contains only reads with variant bases, but no reads with a reference base (i.e., no heterogeneity in that site). There is one site in the ITS sequence of *C. bombacis* ATCC 22867 (accession KT984947), two sites in the ITS sequence of *C. americanus* SA252w (accession OQ540463), and one site in the ITS sequence of *C. africanus* KRAM F-57385 (accession OQ540429) that were incorrectly sequenced.

**Table 4.** Summary of conflicting nucleotide variants of *Ceraceosorus* ITS regions across different detection methods. Details on the conflicting variants can be found in Table S2.

| Conflicting patterns | Possible source of conflict | Numbers of sites/cases with conflicting variant patterns* |  |  |  |  |  |
| --- | --- | --- | --- | --- | --- | --- | --- |
|  |  | ATCC 22867 | MCA4658 | SA252w<sup>a</sup> | KRAM F-57386<sup>a</sup> | KRAM F-57385<sup>ab</sup> | KRAM F-57387<sup>ab</sup> |
| Variant detected in Cloning-Sanger sequencing but not in other methods | PCR-cloning-sequencing error | 9 | 20 | 12 | 10 | 6 | 9 |
|  | 3' end of the amplicon, variant hardly detected in targeted amplicon sequencing | 6 | 3 | 0 | 1 | 1 | 1 |
|  | Ambiguous to identify the source of conflict | 0 | 0 | 0 | 2 | 3 | 4 |
| Variant detected in targeted amplicon sequencing and WGS sequencing, but not in Cloning-Sanger sequencing | Inadequate clone sampling effort | 1 | 0 | 0 | 3 | 1 | 0 |
| Variant falsely called in targeted amplicon sequencing | Error in high-throughput sequencing and read mapping | 5 | 1 | 3 | 0 | 0 | 0 |
| Inconsistent variant calling among independent runs of targeted amplicon sequencing | Inconsistency in high-throughput sequencing | 2 | 2 | 6 | 4 | 0 | 0 |
| Only variant base detected, but not reference base | Inaccurate reference sequence | 1 | 0 | 2 | 1 | 1 | 0 |
| Others |  | 0 | 4 | 0 | 0 | 3 | 0 |
| Total |  | 23 | 28 | 23 | 20 | 15 | 14 |

\*Some cases/sites may have more than one conflicting pattern.
<sup>a</sup> Samples do not have data from WGS sequencing.
<sup>b</sup> Each sample has a single run of targeted amplicon sequencing.

Finally, we re-analyzed the p-distance matrices and phylogenetic reconstruction after correcting PCR-cloning-sequencing errors. The maximum pairwise adjusted p-distances of ITS clones in each strain/specimen are as follows (Table S3): 0.025 for *C. bombacis* ATCC 22867, 0.019 for *C. guamensis* MCA4658, 0.023 for *C. americanus* SA252w, 0.017 for *C. africanus* KRAM F-57386, and 0.025 for *C. africanus* KRAM F-57385 and 0.026 for *C. africanus* KRAM F-57387. The maximum adjusted pairwise p-distance found among all *C. africanus* specimens is 0.029, suggesting that the original p-distance is overestimated due to artefactual variant sites from PCR-cloning-sequencing errors. The phylogenetic tree reconstruction of the ITS region after error correction reveals discrete haplotypes of ITS clones in each *Ceraceosorus* strain (Figure S2). According to our data, there are at least 5 ITS haplotypes in *C. guamensis* MCA4658, 7 haplotypes in *C. bombacis* ATCC 22867 and *C. americanus* SA252w, 9 haplotypes in KRAM F-57386, 11 haplotypes in KRAM F-57385, and 15 haplotypes in KRAM F-57387. Taken together, these results suggest that the ITS regions of all *Ceraceosorus* samples do not evolve in a concerted fashion.

### rDNA analyses from WGS data among Ustilaginomycotina species

We examined intragenomic variation of three rDNA regions (partial 18S, ITS, and partial 28S) in twelve fungal species, representing seven orders of Ustilaginomycotina. *Ceraceosorus* is the only sampled genus with intragenomic variation in all rDNA regions (Table 3). In all *Ceraceosorus* species, the ITS region has the highest number of variant sites, followed by the partial 28S region and the partial 18S region. *Ceraceosorus africanus* is an exception in that the lowest number of variant sites is in the partial 28S region. Meanwhile, only three other species in Ustilaginomycotina contain nucleotide variants in the rDNA regions that are not as highly variable as seen in *Ceraceosorus* (Tables 3, S2). These species are as follows: *Pseudomicrostroma glucosiphilum* MCA4718 with one substitution site in the ITS region (detected by both targeted amplicon and WGS sequencing), *Meira miltonrushii* MCA3882 with one indel site in the partial 18S region (detected only by targeted amplicon Illumina sequencing), and *Tilletiopsis washingtonensis* MCA4186 with one substitution site in the partial 18S region (detected only by targeted amplicon Illumina sequencing).

Although intragenomic variation is absent in most of the Ustilaginomycotina species analyzed in this study, we noticed several nucleotide variants that are falsely called by bioinformatic analyses (Table S2). There are two main reasons for false calling. The first reason is inaccurate reference sequences, even from the same fungal strains/isolates used for targeted amplicon Illumina sequencing and/or WGS Illumina sequencing. This occurs in the ITS and partial 18S regions of *C. guamensis* MCA4658 (accessions: KT984939 and KT984925), the ITS and partial 28S regions of *M. miltonrushii* MCA3882 (accessions NR120190 and JX432962), the partial 18S region of *T. anomala* UBC591 (sequence accession D83193), and the ITS region of *T. washingtonensis* MCA4186 (sequence accession OQ540478, which was sequenced in our study). The second reason comes from errors in false read mapping, which can be noticed from low coverage with read depth less than 100. These false calls are a warning that nucleotide variants retrieved from the bioinformatic pipeline need to be carefully examined and interpreted.

Finally, we estimated rDNA copy numbers in all Ustilaginomycotina species for which WGS data and reference genomes are available. Based on our calculation, the rDNA copy number ranged from 7 to 233 copies with a median of 20.5 (Table 3). The two *Ceraceosorus* species for which genomic data are available have the lowest copy numbers: 7 copies for *C. guamensis* and 8 copies for *C. bombacis*. Two species with the highest estimated rDNA copy number are *T. anomala* UBC591 (233 copies) and *V. palustris* SA807 (114 copies).

## Discussion

### Prevalence of intragenomic rDNA variation in fungi

Our previous work shows high intragenomic variation of the ITS region in two *Ceraceosorus* species—*C. bombacis* and *C. guamensis* (Kijpornyongpan and Aime, 2016). For this study, we expanded our investigation in terms of examined rDNA regions (partial 18S, ITS, and 28S) and taxonomic range (two additional *Ceraceosorus* species, plus representative ‘smut’ fungi for 7 orders of Ustilaginomycotina). As the first subphylum-wide survey of intragenomic rDNA variation in Ustilaginomycotina, we found that only species in the genus *Ceraceosorus* (Ceraceosorales) have a high degree of ITS sequence heterogeneity, which causes failure in obtaining consensus sequences from direct Sanger sequencing attempts (Kijpornyongpan and Aime 2016; Piątek et al. 2016). This failure to directly sequence the ITS region by the Sanger method without ambiguous bases also occurs in several fungi (Pringle et al. 2000; Wang and Yao 2005; Connell et al. 2010; Woo et al. 2010; Alper et al. 2011; Harrington et al. 2014; Liu et al. 2015; Zhao et al. 2015; Roscini et al. 2018). In-depth investigations of many studies also show that intragenomic variation of the rDNA repeats exists in several fungal lineages including Agaricomycotina (Ko and Jung 2002; Wang and Yao 2005; Smith et al. 2007; Lindner and Banik 2011; Vydryakova et al. 2012; Lindner et al. 2013; Cruz et al. 2014), Glomeromycotina (Redecker et al. 1999; Thiéry et al. 2012; Lin et al. 2014; Thiéry et al. 2016; de Souza et al. 2018), Mucoromycota (Woo et al. 2010), Pezizomycotina (Rooney and Ward 2005; Kovács et al. 2011; Li et al. 2013; Naidoo et al. 2013; Poczai et al. 2015; Li et al. 2017; Stadler et al. 2020; Bradshaw et al. 2023), Pucciniomycotina (Moricca et al. 1996, McTaggart and Aime 2018), and Saccharomycotina (James et al. 2009; Dakal et al. 2016; Sipiczki et al. 2018; Colabella et al. 2021). From these studies, we present three aspects of intragenomic variation that need to be considered: 1) variant sites (number of variant sites, from one to dozens of sites; types of variant sites, substitutions, indels or long insertions/deletions), 2) haplotypes (number of haplotypes, from two to twentyish haplotypes; *p*-distance, from < 1% to 20% differences), and 3) frequencies of each variant site/haplotype in the genome (from one count of clone/sequencing read up to 50% frequency). Depending on methodology and data presentation, some studies report on all three aspects, while others report on fewer aspects.

### rDNA sequence heterogeneity and concerted evolution

The concerted evolution theory says that gene homogenization occurs through DNA recombination among rDNA repeats, and this mechanism can repair or remove variant repeats from the consensus sequences of the rDNA repeats (Liao 1999; Eickbush and Eickbush 2007). Because the rDNA repeat in *Ceraceosorus* species appears to have escaped from concerted evolution, we conducted additional analyses to search for potential mechanisms that disable gene homogenization, using genomes generated in previous studies. These are from two *Ceraceosorus* species—*C. bombacis* and *C. guamensis* (Sharma et al. 2015; Kijpornyongpan et al. 2018) and seven other representatives of Ustilaginomycotina (Kijpornyongpan et al. 2018), which appear to fit the concerted evolution model. The genomes of the two *Ceraceosorus* species have the lowest estimated rDNA copy number of all Ustilaginomycotina species analyzed (Table 3). Under the gene homogenization model, we postulate that a homogenization becomes difficult in genomes with lower copy numbers of rDNA repeats. West et al. (2014) found an association between rDNA repeat types in *Saccharomyces cerevisiae* genomes (homogeneous/mosaic) and rDNA copy number. However, analyses of data from Hypoxylaceae species showed no association between rDNA copy number and a number of variant sites (Stadler et al. 2020). Loss of molecular machinery in DNA recombination and repair is another probable mechanism that impedes gene homogenization (Liao 1999; Kobayashi and Sasaki 2017; Paloi et al. 2022). We examined genes that are absent in the *C. guamensis* genome but present in other Ustilaginomycotina genomes. Although we could not attribute any of these missing genes to DNA recombination and repair (Figure S3), further investigation of mutated genes (Kobayashi and Sasaki 2017) in *Ceraceosorus* genomes is promising for a better understanding of an underlying mechanism of concerted evolution.

Although recent studies suggest that intragenomic variation of rDNA regions is widespread in fungi (Stadler et al. 2020; Paloi et al. 2022; Bradshaw et al. 2023), concerted evolution generally maintains sequence homogeneity of rDNA regions. We propose several ways in which rDNA sequences may escape concerted evolution and propagate intragenomic variation. Hybridization is the most common source of rDNA sequence heterogeneity, often found in fungi that are predominantly diploid or dikaryotic (Moricca et al. 1996; Ko and Jung 2002; Wang and Yao 2005; James et al. 2009; Huang et al. 2010; Woo et al. 2010; Vydryakova et al. 2012; Hughes et al. 2013; Naidoo et al. 2013; West et al. 2014; Dakal et al. 2016; Sipiczki et al. 2018; Tremble et al. 2020; Paloi et al. 2022). Naidoo et al. (2013) demonstrated that these heterogeneous rDNA copies can gradually become homogenized after a few generations of sexual reproduction, a process that involves meiosis, crossing-over, and DNA recombination. However, the data from Dakal et al. (2016) suggest that partial homogenization can occur, which creates more variability when considering intragenomic variation in different parts of the rDNA array. Hybridization and partial gene homogenization consequently lead to gene genealogy discordance and the emergence of ‘species complexes’, which are problematic for fungal species identification (Paloi et al. 2022).

The second source of sequence heterogeneity comes from accumulated mutations (Rooney and Ward 2005; Wang and Yao 2005; Lindner and Banik 2011; Li et al. 2013; Lindner et al. 2013; Cruz et al. 2014; Lin et al. 2014; Poczai et al. 2015; Kijpornyongpan and Aime 2016; Li et al. 2017). Mutant rDNA copies can avoid conversion to the original copy (i.e., relaxed concerted evolution) through prolonged asexuality (Lin et al. 2014), ineffective DNA recombination and repair (Kobayashi and Sasaki 2017), or conversion to pseudogenes (Li et al. 2013; Poczai et al. 2015; Li et al. 2017; Sipiczki et al. 2018; Stadler et al. 2020). While subtle rDNA sequence heterogeneity (> 97% similarity) is not a problem for species identification, the emergence of pseudogenes may be, as these may have much lower similarity (ca. 90%) compared to consensus rDNA sequences (Stadler et al. 2020).

The final source of sequence heterogeneity is the introduction of foreign elements, such as group I introns (Bradshaw et al. 2020) or rDNA sequences from distantly related species (Redecker et al. 1999; Virtudazo et al. 2001). These foreign elements can cause a problem not only in species identification, but also in the inaccuracy of genome assembly (Bradshaw et al. 2023). Based on *p*-distance and ITS gene phylogeny, the observed intragenomic variation of *Ceraceosorus* is consistent with an interpretation of accumulated mutation under relaxed concerted evolution, since all ITS haplotypes of each species do not cross species boundaries (Figures 3 and S2, Table S3). Our proposed model for the emergence of intragenomic variation in rDNA is summarized in Figure 4.

**Figure 4.**
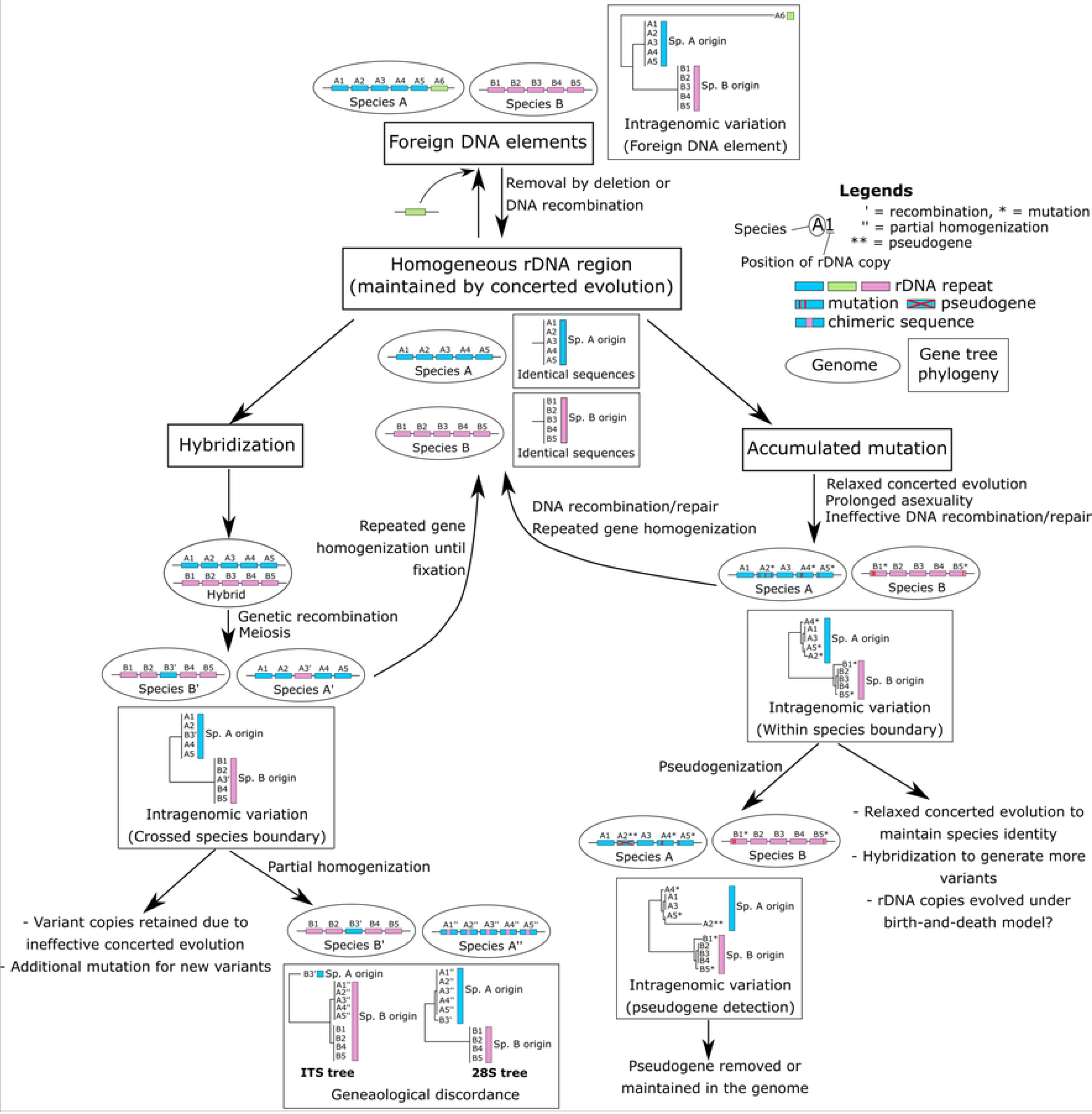
Schematic illustration of possible sources of intragenomic rDNA variation. The model assumes the homogeneity of rDNA sequences by default. We propose three major sources of sequence heterogeneity as follows: hybridization, accumulated mutation, and foreign DNA elements. The rDNA copies in each species/genome are shown in ovals. An alphabet followed by a number indicates the ID for each rDNA copy. Graphs in rectangles illustrate how gene trees look like in each scenario. Branch lengths are proportional to phylogenetic distances. Gene duplications, deletions, and translocations are not shown here to simplify the model.

### Assessment of methods used in detecting intragenomic rDNA variation

Currently, there are several sequence-based methods for detecting intragenomic variation of the rDNA regions in fungi: PCR-direct Sanger sequencing, PCR-cloning-Sanger sequencing, targeted amplicon high-throughput sequencing, WGS high-throughput sequencing, and genome data mining (Paloi et al. 2022). Each method has advantages and disadvantages, which are summarized in Table 5. For instance, PCR-cloning-Sanger sequencing is a traditional method that can provide information about variant sites, haplotypes, *p*-distance, and frequencies of variant sites/haplotypes (depending on the number of clones). However, this method is the largest source of artefacts, which are not found in the other methods (Tables 4, S2). Pairwise *p-*distance values are also significantly reduced after data correction by discarding variant sites found in only one clone (Tables S1, S3). Few previous studies have attempted to deal with PCR errors, such as using high fidelity DNA polymerase (Kovács et al. 2011; Poczai et al. 2015), performing parallel PCR (Kovács et al. 2011), testing secondary clones to check for *Taq* DNA polymerase misreading (Simon and Weiß 2008). However, errors can occur at any stage where a nucleotide base is incorporated into a new strand of DNA. This includes not only PCR amplification, but also multiplication of a cloned vector during bacterial colony growth and the dideoxy-chain termination process during Sanger sequencing. Given this, there is a possibility that several previous studies may have overestimated variant sites, haplotypes and/or *p*-distances, especially those that are represented by data from one sampled clone (Ganley and Kobayashi 2007; Simon and Weiß 2008; Pannecoucque and Hofte 2009; Alper et al. 2011; Thiéry et al. 2012; Vydryakova et al. 2012; Harrington et al. 2014; Liu et al. 2015).

**Table 5.**
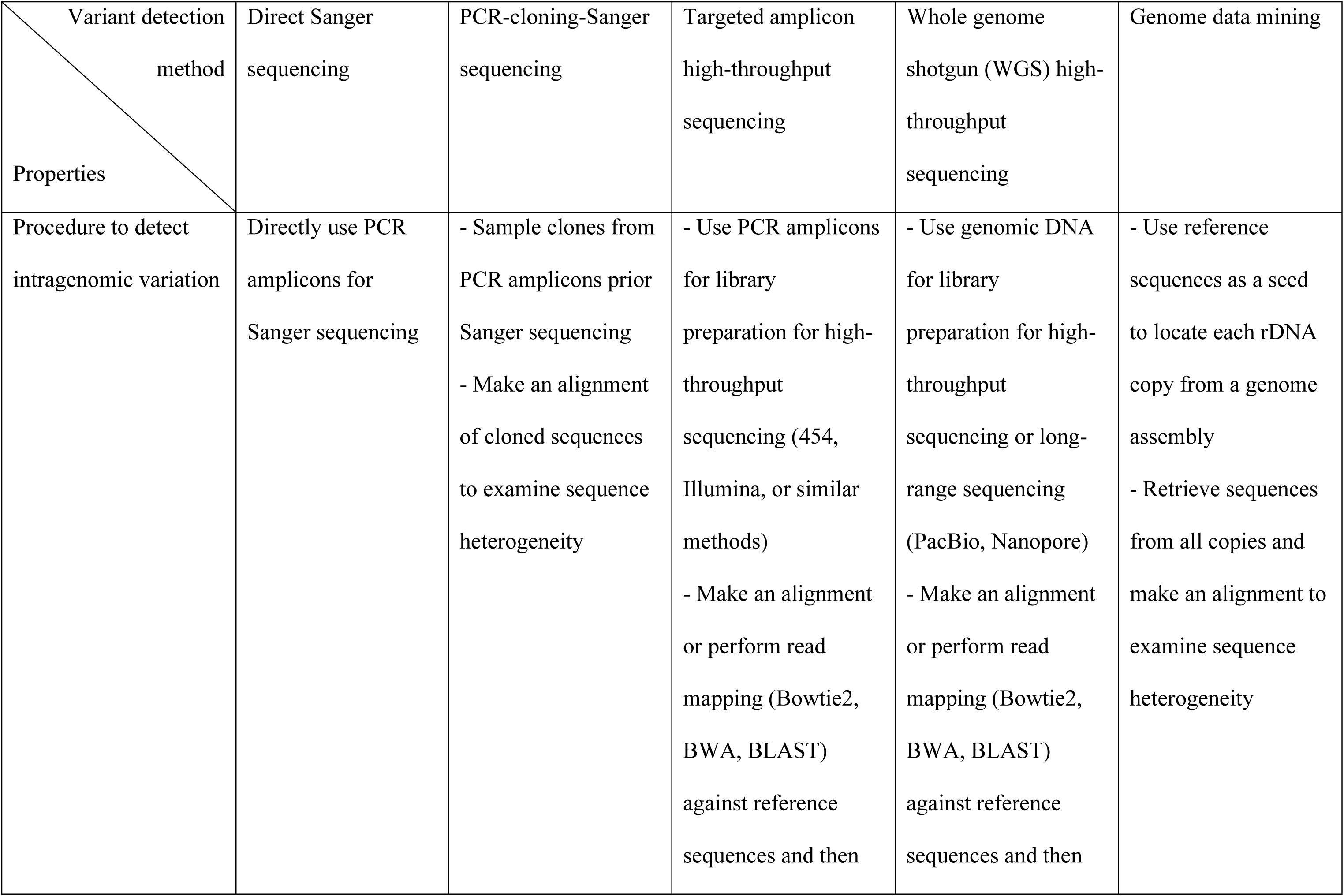
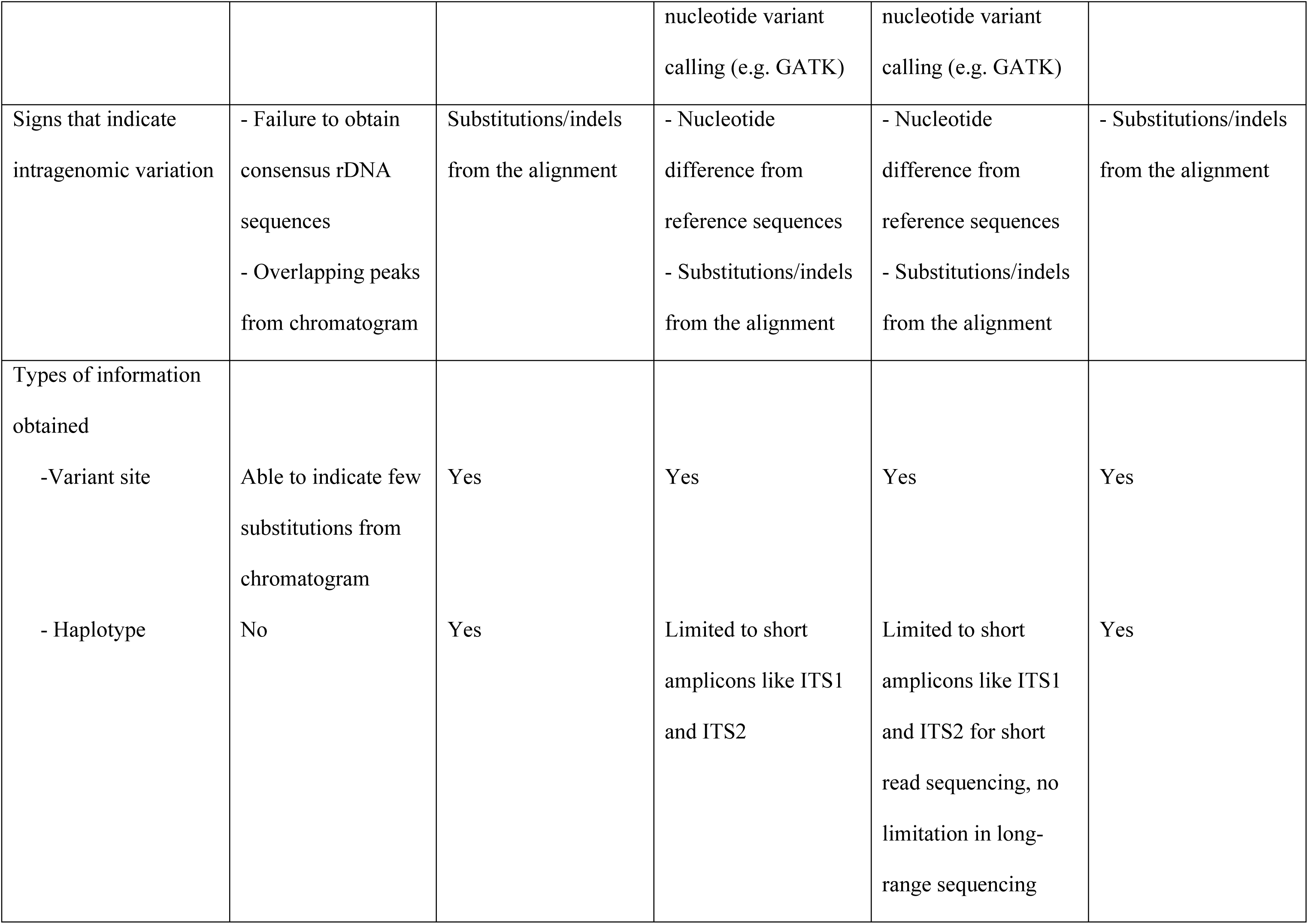
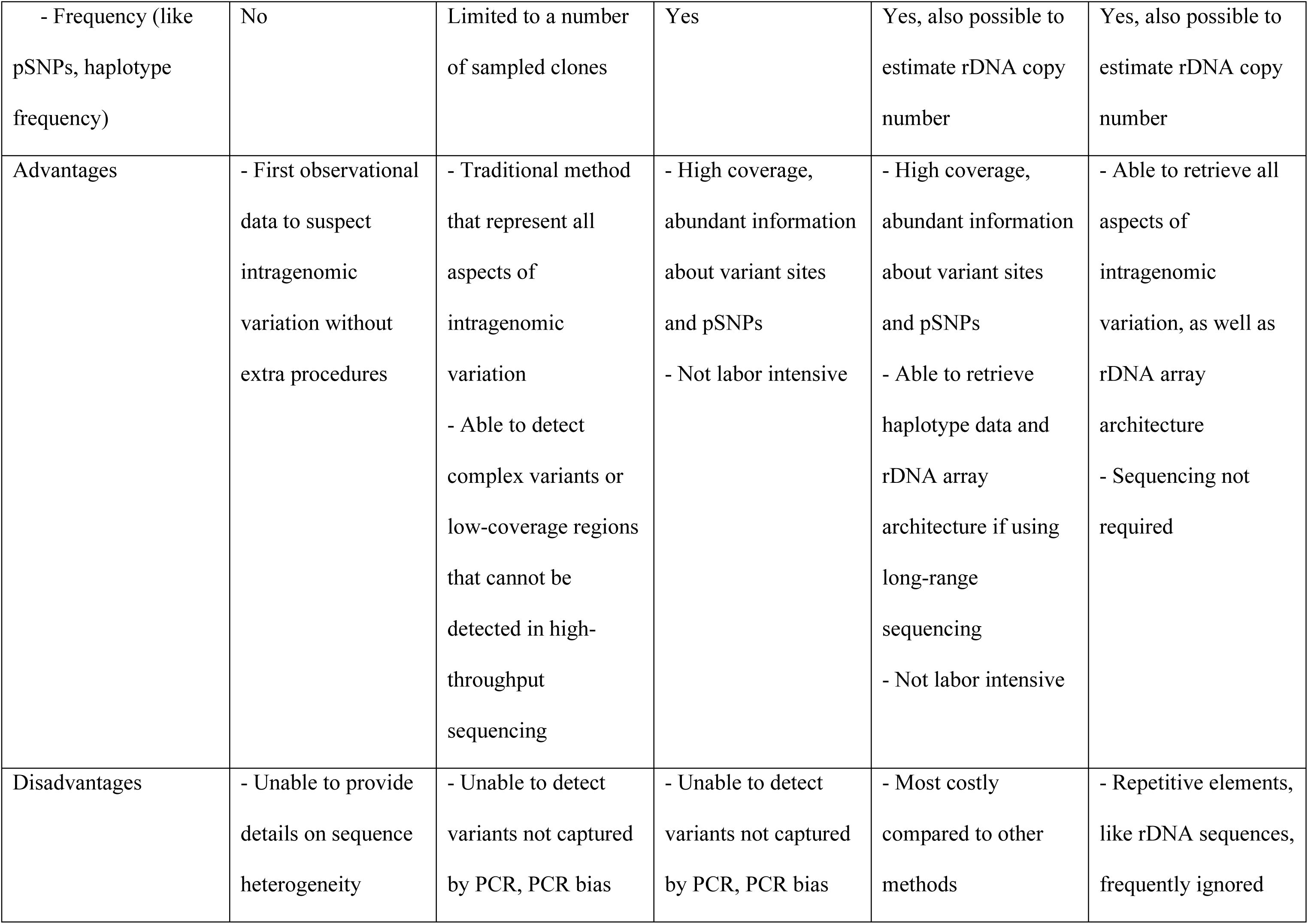
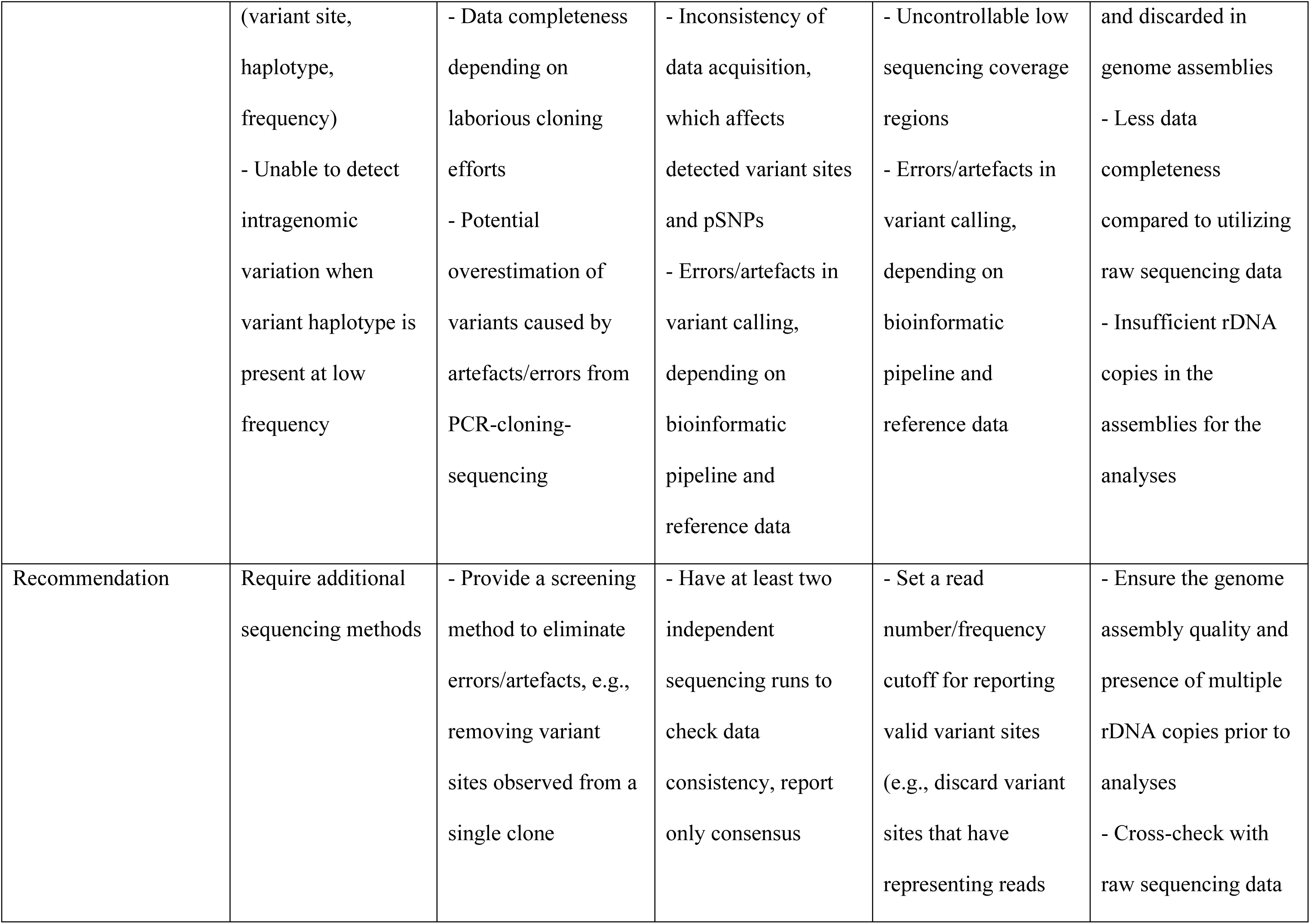
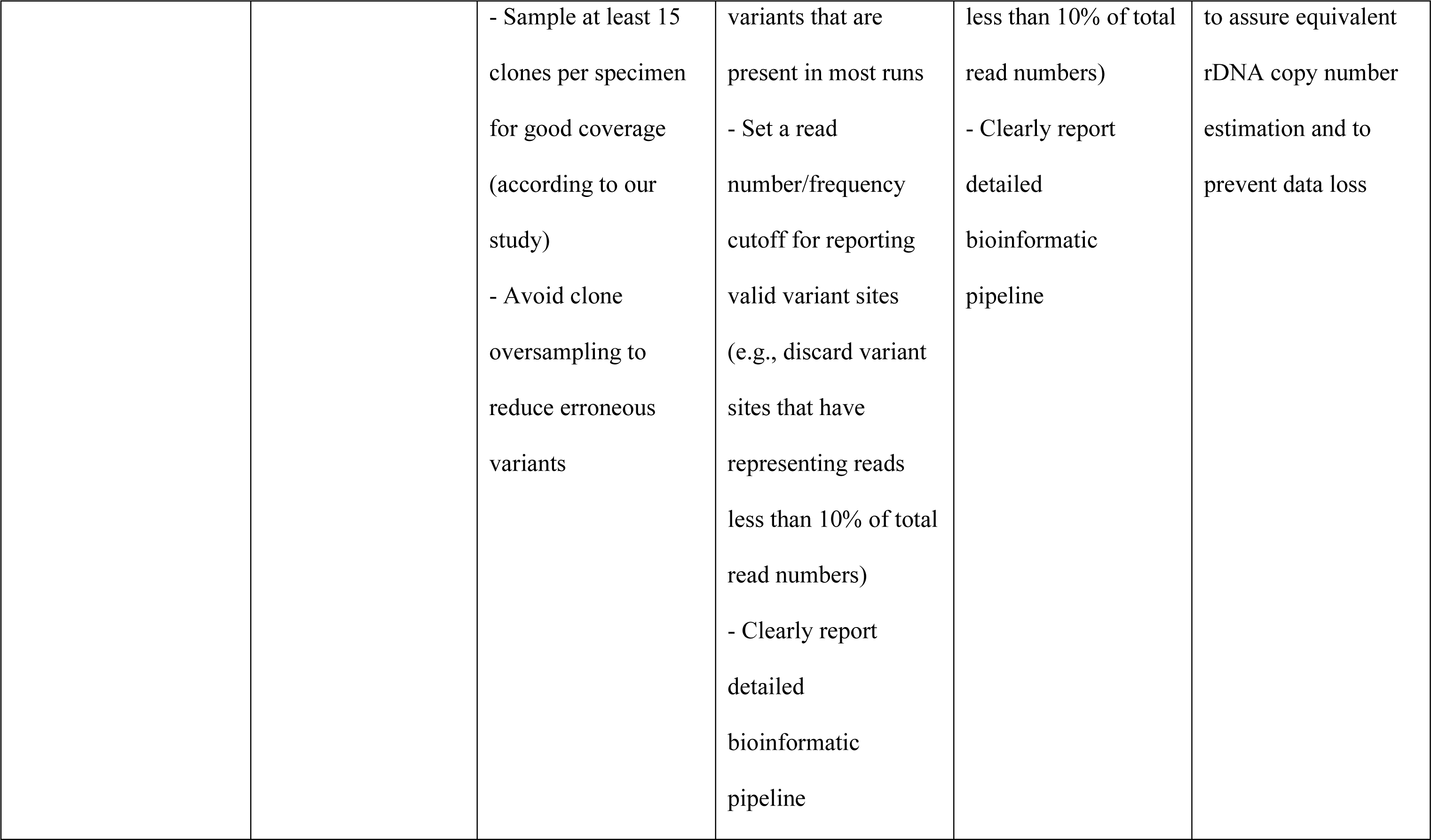
Summarized properties of various detection methods for studying intragenomic variation.

High-throughput sequencing, such as 454 pyrosequencing and Illumina sequencing, is a robust method for generating high sequencing depth and coverage, which allows the detection of rare variant sites not detected by cloning-Sanger sequencing (Tables 4, S2; Lindner et al. 2013; Colabella et al. 2021). This method also allows for the genome-scale study of variant frequencies, such as the recently coined term ‘partial single nucleotide polymorphism (pSNP)’ (James et al. 2009; West et al. 2014). However, there are a few limitations to this method that need to be carefully considered. First, there are two approaches to perform high-throughput sequencing: the targeted amplicon approach and the WGS approach. The former approach may introduce PCR-bias (Tedersoo et al. 2015), especially if rDNA sequences, even functional or pseudogenic, have evolved to avoid a capture by PCR primers (Li et al. 2013; Kijpornyongpan and Aime 2016; Li et al. 2017). Although the latter approach can capture sequences that may be missed during PCR amplification (Lin et al. 2014; this study), it requires high sequencing depth and additional DNA parsing to retrieve sequences of interest. The second limitation comes from data collection. We notice inconsistency in sequence coverage between independent targeted amplicon Illumina sequencing runs on the same sample. This affects the detection of some variant sites, as well as the frequencies of many detected variant sites (Tables 4, S2). The sequencing inconsistency is also found in 454 pyrosequencing (Lücking et al. 2014) and WGS Illumina sequencing (Lin et al. 2014). In addition, sequencing errors in 454 pyrosequencing can lead to many artefactual variant calls (Lücking et al. 2014). The final limitation comes from data analysis. Incorrect read mapping, indicated by low coverage (< 500 mapped Illumina reads), can lead to incorrect variant calls (Tables 4, S2). Stringent read mapping software such as Bowtie2 and BWA may not detect sequencing reads that are too divergent from reference sequences, such as rRNA pseudogenes (Li et al. 2017). Finally, data from short sequencing reads have a limited capacity to perform *p*-distance calculation and haplotype analysis on longer regions such as 18S and 28S rDNA regions.

Genome data mining is a discovery method that can provide information on variant sites, haplotypes, *p*-distances, frequencies of variant sites/haplotypes, and estimated rDNA copy number (Li et al. 2017; Stadler et al. 2020; Paloi et al. 2022; Bradshaw et al. 2023). Despite the potential to analyze all aspects of intragenomic variation within a single platform, the use of genome assembly data requires special precautions. The rDNA region is a part of repetitive elements that are often discarded from the final DNA assembly constructed from short read high-throughput sequencing (Treangen and Salzberg 2012). Thus, in many genome assemblies, the rDNA region is absent or present at much lower copy numbers than in real genomes. A recent study by Bradshaw et al. (2023) reveals that 2251 out of 2414 genomes analyzed have ITS copy numbers < 10, and 694 of them have no ITS sequences present in the assembly. However, Lofgren et al. (2019) estimate that the rDNA copy number of fungal species analyzed in their study ranges from 14 to 1,442 copies. Although genome assemblies constructed from long-ranged sequencing, such as PacBio and Oxford nanopore, tend to have more ITS copy numbers than genome assemblies constructed from short read sequencing (Bradshaw et al. 2023), analyzing data from WGS raw sequencing reads will provide more completeness by preventing information from being discarded during DNA assembly.

Considering our results and many studies from the literature, we provide some recommendations for future research on intragenomic variation of rDNA, as well as similar multigene families. First, having at least two independent runs (i.e., technical replicates) for each type of sequencing is a good approach to avoid inconsistencies in downstream analyses. This is not only to address data inconsistencies from high-throughput sequencing, but also to ensure the accuracy of reference sequences used for nucleotide variant discovery (Tables 4, S2). Second, screening steps are required to remove noise and artefacts from discovered variants. Good practices include discarding or masking variant sites observed from a single clone (Ganley and Kobayashi 2007; Lindner and Banik 2011), setting a minimum frequency cutoff for calling valid variant sites (Colabella et al. 2021), and integrating data from multiple methods. Finally, detailed methods for variant detection must be clearly stated, as they affect the validity of reported variant data. We recommend reporting all aspects of intragenomic variation (as discussed above) whenever possible. Having a standard convention for reporting sequence heterogeneity will be valuable for further comparative analyses across studies to expand the breadth of knowledge of fungal rDNA evolution.

## Conclusion

To date, this study represents the first report of *Ceraceosorus* from the Western Hemisphere. The other species of the genus have been discovered in distant locations in subtropical zones (Figure 2G): *C. africanus* in West Africa, *C. bombacis* in India, and *C. guamensis* in Guam (Cunningham et al. 1976; Kijpornyongpan and Aime 2016; Piątek et al. 2016). Despite their limited number of species and distribution, all examined *Ceraceosorus* isolates/specimens show extensive intragenomic variation, which is not observed in other Ustilaginomycotina lineages. Therefore, the genus *Ceraceosorus* can serve as a promising model to study relaxed concerted evolution in the rDNA region through comparative genomics and experimental genetics. We anticipate that our results will have implications for future research in taxonomy, systematics, metagenomics, evolutionary biology, genomics, and any other disciplines that uses rDNA as a tool for study.

## Data Availability Statement

Reference rDNA sequences and ITS clone sequences generated from this study are available in the NCBI GenBank database under the accessions provided in Table 1. WGS sequencing data used in this study were retrieved from the DOE-Joint Genome Institute (JGI) Fungal Genomics Program and will be available upon request. Other targeted amplicon high-throughput sequencing data, including bioinformatic scripts used in the analysis are available in the DRYAD repository (https://doi.org/10.5061/dryad.qbzkh18r3). The temporary link to access the unpublished datasets during a peer-review process is https://datadryad.org/stash/share/cd4rk2pWiYFahE0-Lg2KQTQz2AicP9VOv9s2dDqrbLU.

## Acknowledgement

We would like to thank Dr. Phillip SanMiguel and staff from the Purdue Genomics Core Facility for assistance in high-throughput sequencing. This article is in memory of Rick Westerman, a former staff member of Purdue Genomics Core who passed away in September 2020. He provided significant valuable advice on data analyses for our study. This research was funded in part by the US National Science Foundation [grant number DEB-2127290] and the USDA National Institute of Food and Agriculture Hatch project [grant number 1010662].

